# Transcriptome profiling of *Variovorax paradoxus* EPS under different growth conditions reveals regulatory and structural novelty in biofilm formation

**DOI:** 10.1101/2019.12.17.879619

**Authors:** Richard J. Fredendall, Jenny L. Stone, Michael J. Pehl, Paul M. Orwin

## Abstract

We used transcriptome analysis by paired-end strand specific RNA-seq to evaluate the specific changes in gene expression associated with the transition to static biofilm growth in the rhizosphere plant growth promoting bacterium *Variovorax paradoxus* EPS. Triplicate biological samples of exponential growth, stationary phase, and static biofilm samples were examined. DESeq2 and Rockhopper were used to identify robust and widespread shifts in gene expression the transcriptomic signals specific to each growth phase. Weidentified 1711 protein coding genes (28%) using DESeq2 that had altered expression greater than 2-fold specifically in biofilms compared to exponential growth. Fewer genes were specifically differentially expressed in stationary phase culture (757, 12%). A small set of genes (103/6020) were differentially expressed in opposing fashions in biofilm and stationary phase, indicating potentially substantial shifts in phenotype. Gene Ontology analysis showed that the only class of genes specifically upregulated in biofilms were associated with nutrient transport, highlighting the importance of nutrient uptake in the biofilm. The biofilm specific genes did not overlap substantially with the loci identified by mutagenesis studies, although some were present in both sets. The most highly upregulated biofilm specific gene is predicted to be a part of the RNA degradosome, which indicates that RNA stability is used to regulate the biofilm phenotype. Two small putative proteins, Varpa_0407 and Varpa_3832, are highly expressed specifically in biofilms and are predicted to be secreted DNA binding proteins, that may stabilize extracellular DNA as a component of the biofilm matrix. An flp/tad type IV pilus locus (Varpa_5148-60) is strongly downregulated in specifically in biofilms, in contrast with results from other systems for these pili. Mutagenesis confirms that this locus is important in surface motility rather than biofilm formation. These experimental results suggest that *V. paradoxus* EPS biofilms have substantial regulatory and structural novelty.

## INTRODUCTION

It is well established that many if not most bacteria spend a large fraction of their time in the biofilm state [1]. Many models of biofilm formation and growth have been evaluated, and while there are differences in the outcomes [2], one widely accepted static biofilm model is attachment to an abiotic surface while immersed in growth medium [3]. This model has been used to study biofilm formation in pathogens, plant associated bacteria, and industrial biofouling, and life attached to natural surfaces (for review see [1, 4]). Biofilm gene expression patterns have been examined in many systems – a search of the sequence read archive (SRA) for the terms “transcriptome and biofilm” yields 922 results! The rapid advancement of DNA sequencing and reduction in cost have made genomes and transcriptomes much more widely available beyond the traditional model microorganisms [5]. Transcriptome analysis can lead to insights into an organism’s response to changing physiological conditions that evade traditional mutagenesis approaches [6]. The costs for these analyses have dropped precipitously, and the number of bacterial whole genomes available is very large (74,764 on 12-10-19, https://img.jgi.doe.gov/cgi-bin/m/main.cgi?section=ImgStatsOverview), especially in the easily cultivatable groups, making it possible to use these approaches outside the common model organisms. Shifts in gene expression have been specifically associated with the biofilm lifestyle, across many different bacteria, and in different biofilm contexts [3, 7–9]. A wide distribution of gene expression patterns have been observed, and there is a great deal of variability that is both strain specific and driven by experimental setup.

*Variovorax paradoxus* is a soil dwelling member of the beta-proteobacteria that is widely recognized as an important plant-growth promoting bacterium [10, 11]. It has been frequently identified as a degrader of xenobiotics and source of important enzymes for biocatalysis [10]. Several finished genomes of *Variovorax paradoxus* strains are published [12, 13], and many more are available as permanent draft sequences (twenty annotated genomes including four complete genomes at ncbi.nlm.nih.gov). The strain that we discuss here, *V. paradoxus* EPS, has been evaluated for its behavior in laboratory conditions that regulate swarming motility and biofilm formation [14], and we have also used insertional mutagenesis to identify some genes involved in these complex phenotypes [15]. In this work we show that the shift to biofilm growth in *V. paradoxus* is accompanied by a large-scale change in transcript profile. The biofilm gene expression profile is substantially different from the stationary phase profile, and the robustness of the signal was confirmed using two independent analyses of the raw transcript data. A total of 1711 genes were found to be uniquely and significantly altered in expression by more than 2-fold in the biofilm cultures, representing 28% of protein coding genes (1711/6020). A much smaller number of genes was identified as specifically differentially regulated in stationary phase culture (757/6020, 12%). There were substantial deviations in the number of genes identified using different analysis tools, but most of the deviations were related to *de novo* gene identification and data normalization strategies, and do not significantly alter the overall picture. Shifts previously observed in single gene qPCR are observed again [15], providing further independent confirmation of the consistency of the data. Only a few of the genes previously identified as affecting biofilm formation were present in the significantly altered transcript profiles, and all of the ones identified were among the genes downregulated in biofilm. The most highly upregulated biofilm specific gene, Varpa_1640 encodes a putative DEAD-box helicase that is predicted to be part of the RNA degradosome, implying that biofilm transition may be globally regulated by RNA stability. Two small hypothetical proteins with homology to the SNF2 superfamily of DNA binding proteins are also highly upregulated. Both of these proteins have predicted signal peptides in their sequences that lead to the hypothesis that they are non-specific DNA binding proteins that stabilize the biofilm matrix. In contrast to many other previously described biofilm systems, pili appear to be downregulated, and the transcriptome data on the *tad* locus (Varpa5148-60) along with mutational analysis suggests that this pilus is involved in motility rather than attachment. All of these insights and potential new avenues for exploration come from analysis of the gene expression pattern, and most were not accessible by traditional mutational analysis.

## MATERIALS AND METHODS

### Culture conditions

All cultures were grown in 2.5 g/L yeast extract (50% YE) for 24hrs from a isolated colonies picked off of a low passage plate (1-2 passages since −80 storage). All cultures were incubated at 30**°**C at all times, liquid growth cultures were incubated with shaking at 200 rpm for maximal oxygenation. For logarithmic growth, the culture was diluted 1:20 into 10 mL of 50% YE, and monitored spectrophotometrically using OD595. Aliquots from three separate cultures were collected when the OD595 was approximately 0.5, and RNA was immediately extracted from ~10^9^ cells per sample. Identical cultures were grown for stationary phase analysis, but RNA was collected when the OD595 was stable for successive readings spaced 30m apart (approximately 18h after dilution inoculation). For biofilm cultures, an identical dilution was grown for 24h in 12 well Falcon plates (non-tissue culture treated) with 2 ml of liquid medium per well. The plates were incubated at an angle using a 10 ml serological pipet to create an air/liquid interface on the bottom of the well. After 24h the medium was replaced with minimal disruption to the biofilm, and the culture was incubated for an additional 24h under the same conditions. After this incubation the liquid medium was removed, the plate was washed with fresh medium, and the biofilm was recovered by scraping. One well in the plate was incubated with media only as an inoculum control. The 11 inoculated wells from each plate constituted a biological replicate.

### RNA isolation

Samples were pelleted by centrifugation at 10000 x g and the supernatant was discarded. The pellet was resuspended in 200 μL of 1mg/mL lysozyme in 10 mM Tris 1mM EDTA (TE, pH 8.0). Two volumes of RNAprotect (Qiagen) was added to each sample and the samples were incubated for 10 minutes at 25 °C. Some samples were preserved in this solution at −80 °C until further processing. The remaining steps were performed following the RNeasy (Qiagen) protocol. Successive addition of 700 μL of Buffer RLT and 500 μL of 100% ethanol (RNA grade, Fisher) was followed by transfer of the mixture into an RNeasy Mini Spin column. After centrifugation the flow through was discarded. The column was washed with 350 μL of buffer RW1. To eliminate DNA from the sample, 100 μL of RQ-1 RNase free DNase (Promega) was added to the column and incubated at 25°C for 15 minutes. The column was then washed again with 350 μL of RW1 and the remaining steps of the protocol were followed.

### RNA purification and assessment

RNA was precipitated by adding 10% (v/v) 3M sodium acetate, 5 μg glycogen and 3 volumes of 100% ethanol, and incubating overnight at −20°C. The RNA was pelleted by centrifugation in an Eppendorf 5810R centrifuge using a FA 45-30-11 rotor at 12000 x g for 30 minutes at 4°C. The liquid was carefully by aspiration removed and the pellet resuspended in 1 mL of 70% ethanol. After an additional identical centrifugation step, the pellet was dried and then resuspended in 24 μL of TE. The concentration and purity of the RNA samples was determined spectrophotometrically using a NanoDrop ND1000 spectrophotometer (Thermo Fisher). Further analysis for sample suitability was performed by Beijing Genomics Institute (Hong Kong).

### RNA-seq

The RNA was sequenced using a strand-specific paired end protocol by BGI genomics (Hong Kong). Each sample was sequenced for 91 cycles in an Illumina HiSeq instrument with a total of 1.5×10^7^ reads per sample. Each of the three conditions (logarithmic, stationary, biofilm) was sampled in triplicate (biological replicates), derived from single colonies plated on YE agar directly from the original *V. paradoxus* EPS stock culture. The raw sequences were transmitted as fastq files for subsequent analysis. These files have been uploaded to NCBI and are available as gzip archives (BioProject <u>PRJNA594416</u>, BioSample <u>SAMN13517278,</u> SRA accession #s <u>SRR10613920-8</u>).

### Expression analysis

Raw sequence data was uploaded to the Galaxy main server (galaxy.psu.edu) and all of the bioinformatic tools described below were accessed at that site, unless otherwise noted. Uploaded sequences were trimmed based on quality using the Trimmomatic tool [16], and aligned to the *Variovorax paradoxus* EPS genome using BWA [17]. Trimmed sequences were also aligned to the genome using the Rockhopper suite of RNA-seq tools for differential gene expression analysis [18] on a local desktop computer. The BWA alignment was transformed into count data using the StringTie tool [19], and differential expression analysis was undertaken using the DESeq2 version 1.18.1 toolkit following the workflow outlined in [20]. The DESeq2 program treated each individual gene as a separate transcript for the purposes of differential expression analysis. Venn diagrams were drawn in the Galaxy tool using the output of DESeq2 filtered on >2x change in expression level, and tables of up and down regulated genes were generated using the Filter tool on Benjamini-Hochberg adjusted p-value p <0.05 and fold expression change > 2. Transcript counts from Rockhopper were visualized (.wig files) in the Integrated Genome Viewer (IGV) available at (https://software.broadinstitute.org/software/igv/). Additional Rockhopper text file output on operon structure and *de novo* transcript structure were used for analysis of specific individual genes.

### Motility mutant screen and complementation

Transposon mutants were generated using Tn5 (tetR) as described previously [15]. Mutants with impaired motility were enriched in YE (5 g/L) liquid culture by growth with shaking overnight followed by a 1h settling period, followed by transfer of 1:100 of the culture volume into a fresh tube. This procedure was repeated x times (look at Jenny’s proposal) followed by plating on swarming medium [14]. Isolates that were identified as having decreased motility were subsequently tested for stability of the defect by repeated swarming assays. The interrupted gene was identified by the rescue cloning approach described previously [15]. PCR amplified fragments from the *V. paradoxus* EPS genome corresponding to Varpa_5148 alone and the region including Varpa_5148-50 were cloned including regulatory sequence identified from transcriptome examination into pCR2.1 (KanR) (Invitrogen) and verified by Sanger sequencing. These elements were subcloned into pBBR1MCS-2 (KanR) [21] using *Bam*HI and *Hin*dIII sites incorporated into the primers. The complementation constructs and control plasmids were then introduced into *V. paradoxus* EPS by electroporation as described previously [15].

## RESULTS AND DISCUSSION

### Differential Expression analysis using DESeq2

The DESeq2 suite of programs was used to analyze transcripts from all three growth conditions in a pairwise fashion [20]. All of the outputs of DESeq2 are included in **Supplemental Data S1**. The three sets of samples were evaluated for sample consistency among biological replicates using PCA analysis (Figure 1A) and a heatmap of Euclidian distance (Figure 1B). The three biological replicates from each sample are clustered together as expected, and exponential growth samples (“log”) are less variable than either the stationary phase or biofilm samples. Using the Benjamini-Hochberg false discovery rate, the adjusted p-values were calculated for each pairwise comparison. The MA plots of comparisons between log phase and either stationary phase (Figure 2A) or biofilm (Figure 2B) show the distribution of differential expression. The top 25 up and down regulated loci unique to biofilm growth are listed in Table 2 and **Table 3**. Loci identified by DESeq2 but not corresponding to an open reading frame were ignored for this analysis but included in the global expression analysis. After filtering for >2x change in expression, 1104 loci were identified as having significantly increased expression uniquely during biofilm growth, while 607 had decreased expression using the same parameters (Figure 3A, B). Interestingly, the unique proportion of the differentially regulated genes in each direction (1104/1436 v 607/865) was similar. An additional set of comparisons revealed a total of 103 genes which where expression relative to exponential growth was regulated in opposite directions between biofilm and stationary phase (Figure 3 C, D). The lists of genes specifically differentially expressed in biofilm and stationary phase, and the list of genes with opposite regulation, are listed in **Supplemental Data S1** along with RPKM values and DESeq2 derived significance values.

**Figure 1.**
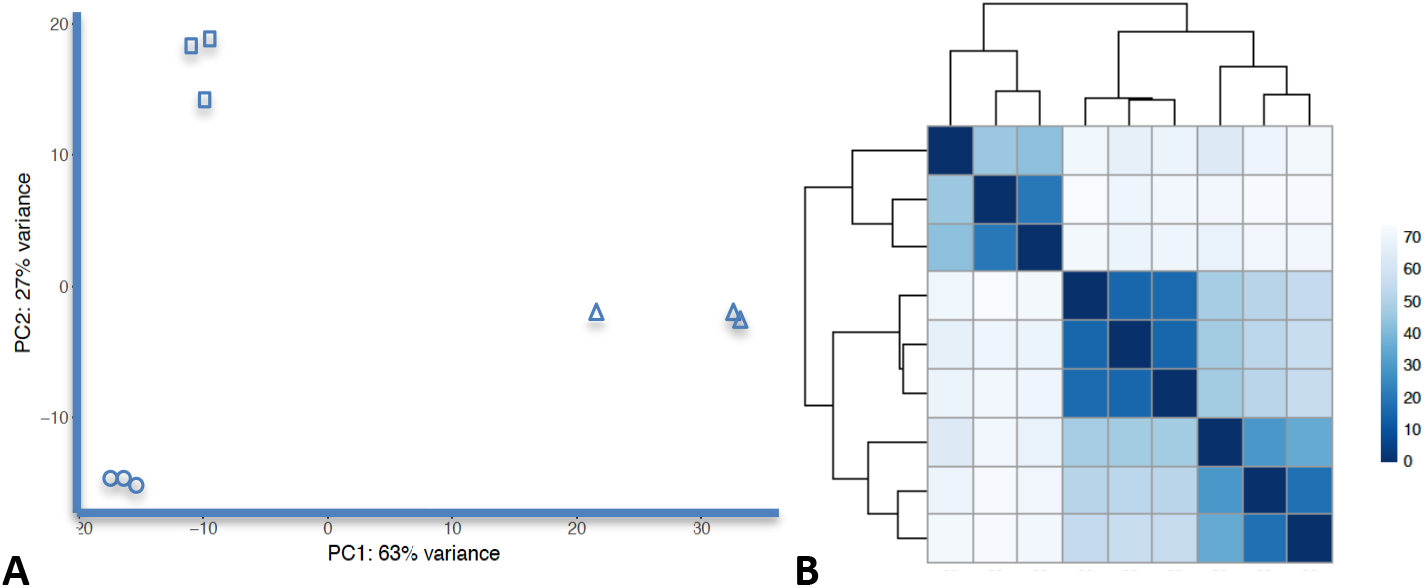
**A**) PCA analysis of datasets using DESeq2 shows clustering of Log phase (circles), stationary phase (squares) and biofilm (triangles) samples. **B**) Heatmap of Euclidian distance shows similar clustering of samples, indicating biological replicate consistency.

**Figure 2.**
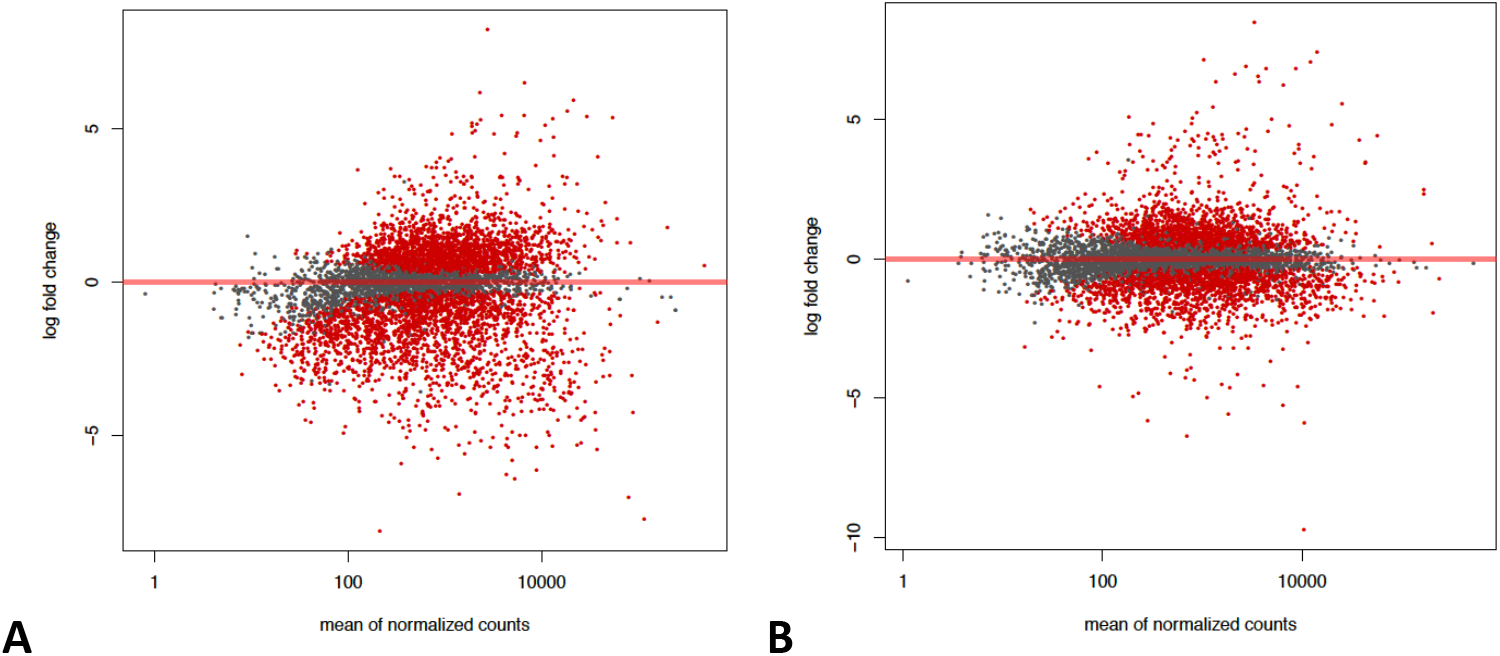
MA plots. A) exponential planktonic growth compared to biofilms B) exponential planktonic growth compared to stationary phase. Red dots indicate adjusted p-value (q-value) of <0.1).

**Figure 3.**
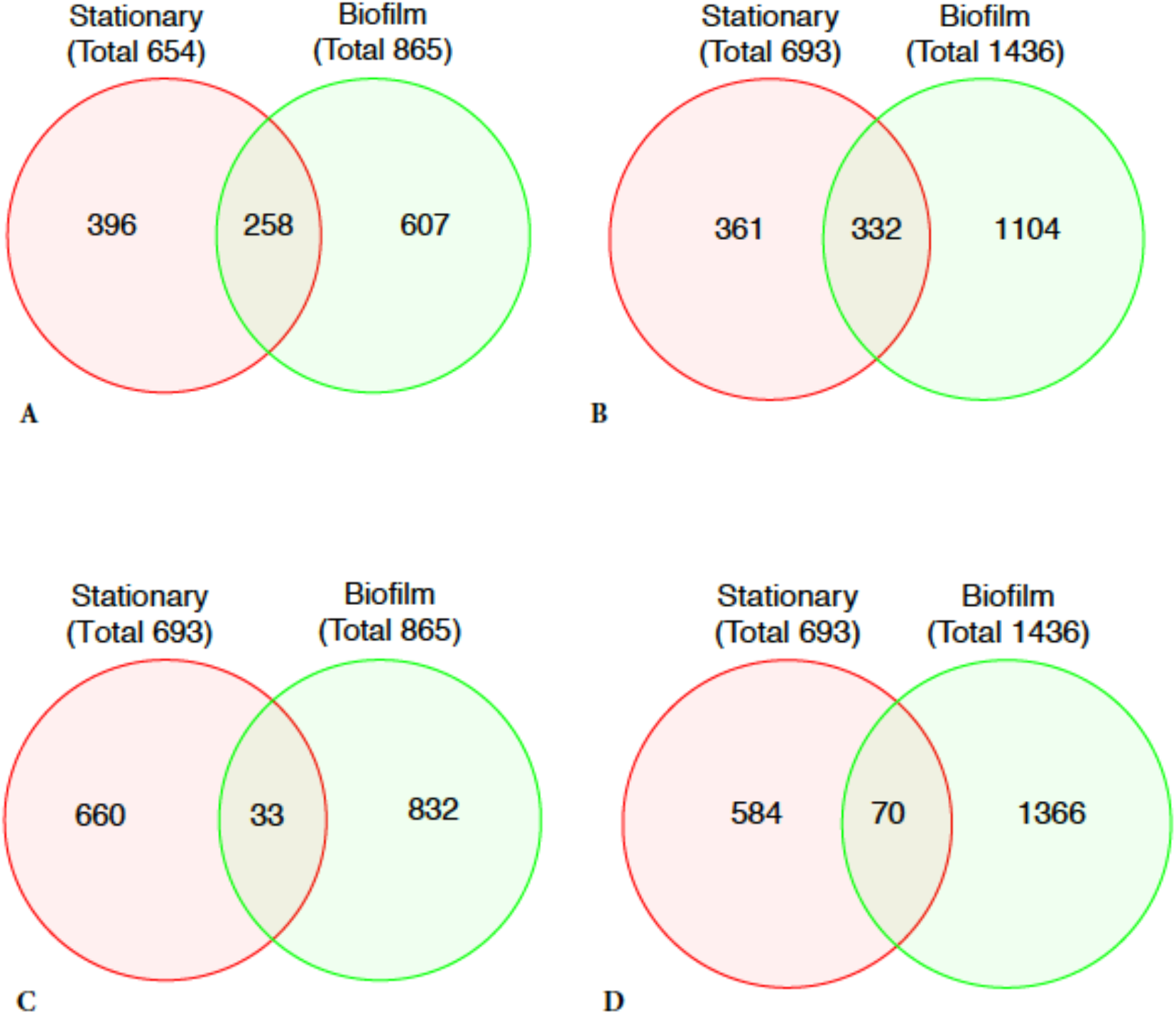
Venn Diagrams of genes differentially expressed in stationary phase or biofilm compared to exponential growth. Output of the DESeq2 analysis pipeline were filtered using 2-fold expression as a cutoff. **A)** Genes downregulated 2-fold or more. **B)** Genes upregulated 2-fold or more. **C)** Genes upregulated in stationary phase and down in Biofilm. **D)** Genes downregulated in stationary phase and upregulated in Biofilm.

### Transcriptome structure analysis with Rockhopper

Pairwise analysis of the transcriptome replicates was performed using Rockhopper [18] to compare the methods and evaluate the robustness of the datasets. The overall results of differential expression analysis were similar (not shown) with differences attributable to count normalization and annotation parameters. We limited our use of Rockhopper to in depth analysis of individual loci and to identification of operon structure. This program will also identify small RNA, but since our initial data collection did not specifically isolate small RNA, data on differential small RNA expression is not presented here. 1262 multigene operons were identified by Rockhopper based on transcript data, along with 3021 gene pairs. The total number of protein coding genes identified as differentially regulated in biofilms was 2145, while the number differentially regulated in stationary phase was 2508. This was a substantially different outcome than with DESeq2. When examining the text outputs of the comparison, it was observed that Rockhopper and DESeq2 both identify many potential small RNAs and other non-coding elements, which are not directly comparable. In all cases where an individual gene was evaluated for differential expression, the results were similar, and quantitative differences in fold-induction or repression are likely due to different normalization or *de novo* transcript identification algorithms. All of these results are included in **Supplemental Data S2**.

### Loci previously evaluated for biofilm transcription

Our previous work identified a number of loci that when mutated resulted in altered biofilm and/or swarming phenotypes [15]. We found that only 8/30 loci identified previously by transposon insertion [15] were differentially regulated at the level of transcription, and all of those loci were downregulated in biofilms. In that work several loci were identified by multiple insertions that were all associated with the phenotypic alteration, namely multiple insertions into Varpa_5900 and Varpa_4680, encoding the PilY1 tip adhesin and a glycosyl transferase, respectively. In both of these cases the RNA-seq analysis confirmed the data collected previously by qRT-PCR [15], confirming the validity of the assay conditions independently. Interestingly, there are 3 PilY1 loci in *V. paradoxus* EPS (5900, 3518, 4912), but only Varpa_5900 is expressed to a significant degree under any of the conditions tested.

### Gene Ontology Enrichment

Gene ontology analysis of the upregulated and downregulated loci is presented in Figure 4. Panels **A** and **B** show diverse sets of gene categories that are overrepresented among the genes with differential expression in biofilms, while panel **C** identifies a much narrower set of genes overrepresented in the set of genes upregulated specifically in biofilms. No gene categories were overrepresented among the loci specifically downregulated in biofilms. The overrepresented categories in Figure 4C are indicative of the increased need for transport of essential micronutrients such as metal co-factors and nutrients that tend to limit replication and viability such as inorganic phosphorus. This latter gene family may also be connected to the use of eDNA as a nutrient storage mechanism as well as a structural component of the biofilm matrix [22]. These GO category patterns are consistent with a biofilm structure with increased nutrient transport to maintain viability throughout the structure and to distribute resources throughout the biofilm.

**Figure 4.**
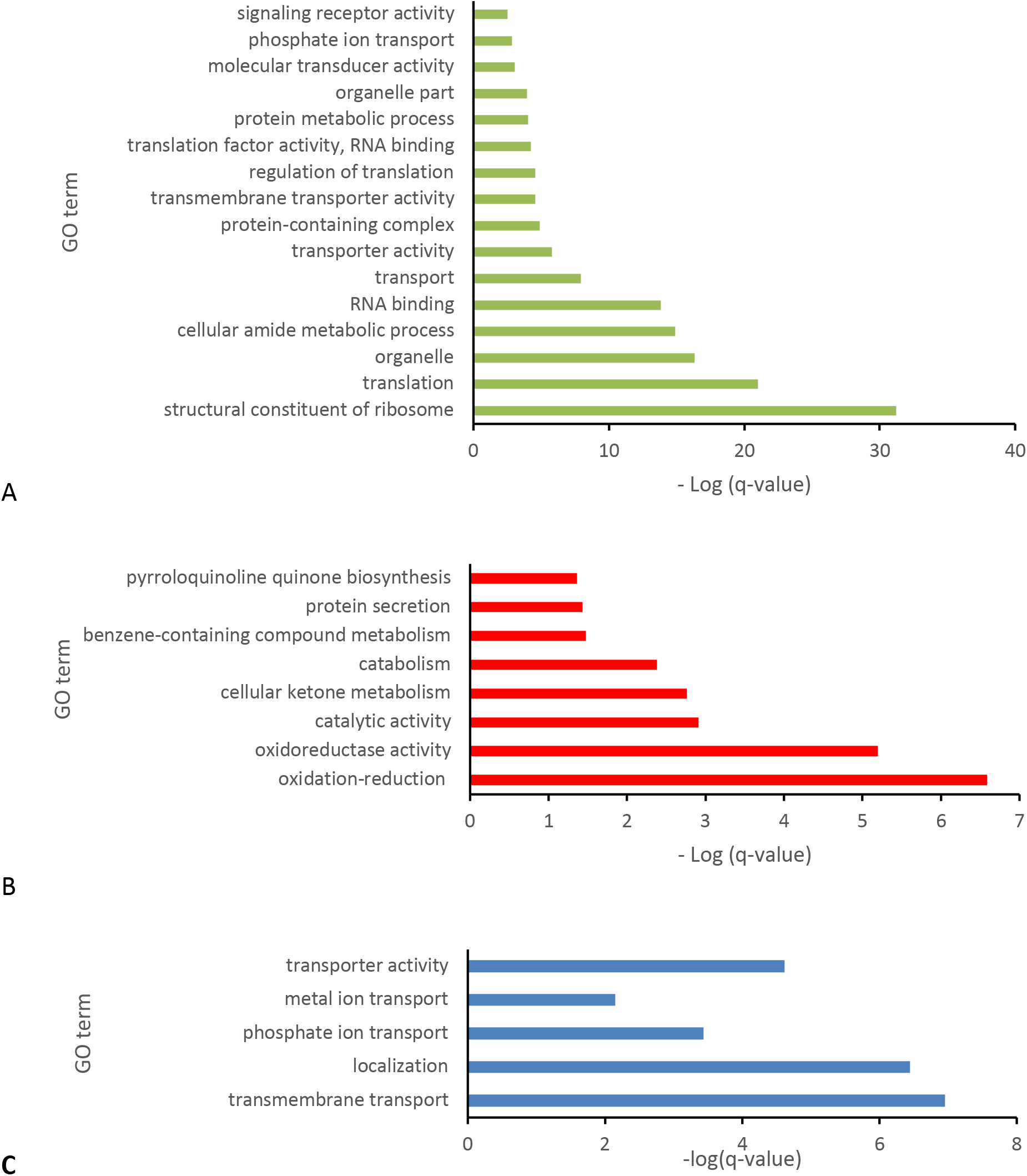
Analysis of Biofilm upregulated gene classes (A) and Biofilm downregulated gene classes (B) compared to whole genome. Panel (C) shows the GO enrichment of genes only upregulated in Biofilm and not in Stationary phase cultures. No GO terms were found to be enriched in genes downregulated only in Biofilms.

### Secreted snf2 family proteins

A pair of small proteins were specifically highly upregulated in biofilm growth (Table 1, **red**), which have high homology to one another (Varpa_0407 and Varpa_3832). Highly similar proteins are found in some other *V. paradoxus* genomes examined, but no close orthologs were found outside the genus using BLASTp [23]. Both genes are highly expressed specifically in the biofilm state, and both have an extensive 5’ untranslated region (Figure 5A, only Varpa_0407 shown). They are annotated as members of the Snf2 superfamily of DNA binding proteins [24, 25], but were much smaller than all previously described proteins in this superfamily. The proteins in *Variovorax paradoxus* are also annotated as having a 24 amino acid leader peptide cleaved in each case to generate a mature 102 amino acid protein. The most common association of Snf2 proteins with function is with helicase activity and chromatin remodeling [25], neither of which is compatible with a bacterial secreted protein. Our conjecture based on this information is that these small proteins are non-specific DNA binding proteins that are secreted to stabilize the biofilm structure which likely contains extracellular DNA (eDNA) as a structural component, as is common in many bacterial species [22]. The presence of these proteins may stabilize the biofilm matrix and protect against extracellular DNAse activity (Figure 5B). It also may be that they were not uncovered by mutational analysis because of their likely functional redundancy.

**Table 1.**
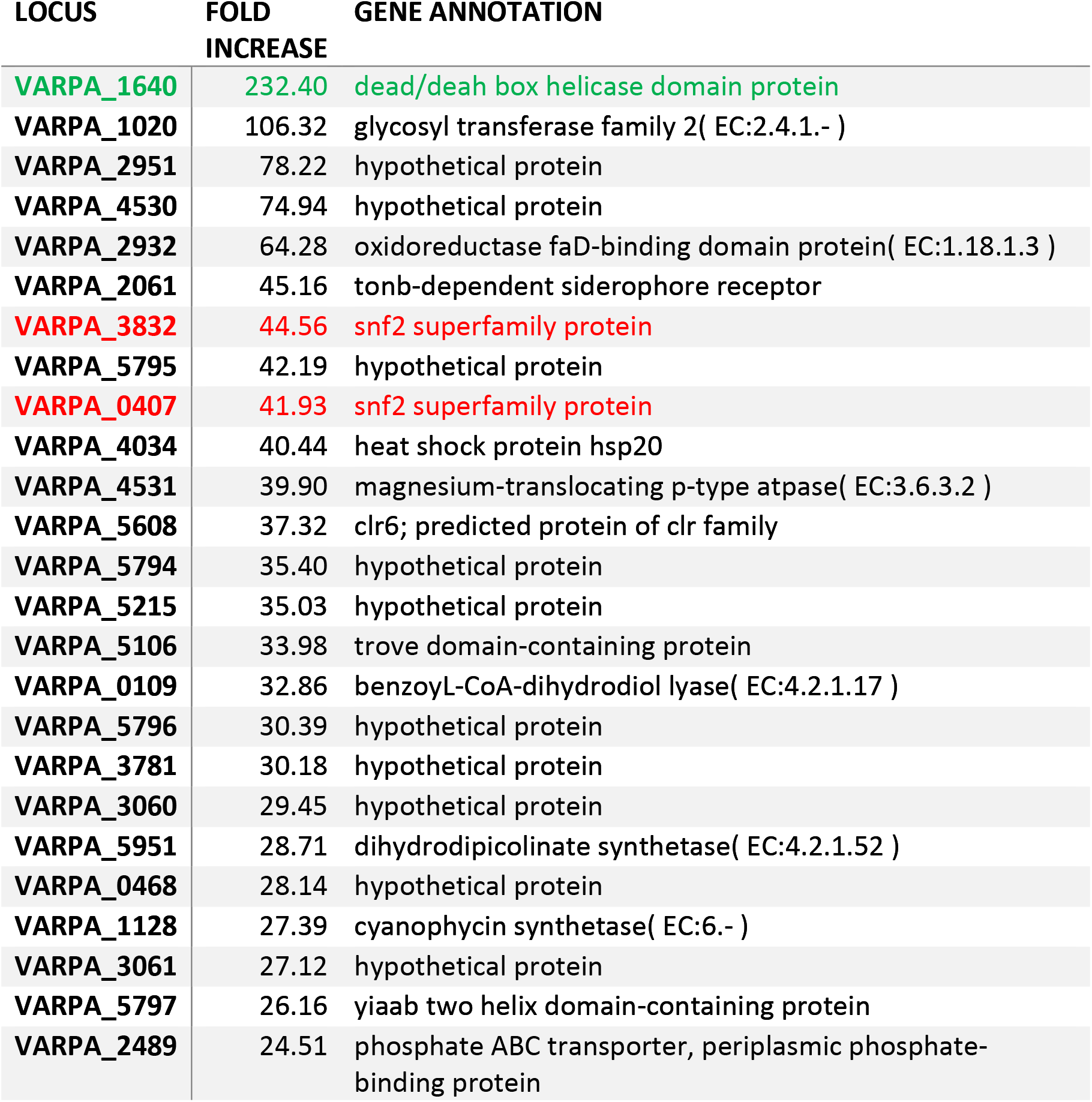
Top 25 uniquely upregulated loci in biofilm cultures

| LOCUS | FOLD<br>INCREASE | GENE ANNOTATION |
| --- | --- | --- |
| **VARPA_1640** | **232.40** | **dead/deah box helicase domain protein** |
| **VARPA_1020** | 106.32 | glycosyl transferase family 2( EC:2.4.1.- ) |
| **VARPA_2951** | 78.22 | hypothetical protein |
| **VARPA_4530** | 74.94 | hypothetical protein |
| **VARPA_2932** | 64.28 | oxidoreductase faD-binding domain protein( EC:1.18.1.3 ) |
| **VARPA_2061** | 45.16 | tonb-dependent siderophore receptor |
| **VARPA_3832** | **44.56** | **snf2 superfamily protein** |
| **VARPA_5795** | 42.19 | hypothetical protein |
| **VARPA_0407** | **41.93** | **snf2 superfamily protein** |
| **VARPA_4034** | 40.44 | heat shock protein hsp20 |
| **VARPA_4531** | 39.90 | magnesium-translocating p-type atpase( EC:3.6.3.2 ) |
| **VARPA_5608** | 37.32 | clr6; predicted protein of clr family |
| **VARPA_5794** | 35.40 | hypothetical protein |
| **VARPA_5215** | 35.03 | hypothetical protein |
| **VARPA_5106** | 33.98 | trove domain-containing protein |
| **VARPA_0109** | 32.86 | benzoyl-CoA-dihydrodiol lyase( EC:4.2.1.17 ) |
| **VARPA_5796** | 30.39 | hypothetical protein |
| **VARPA_3781** | 30.18 | hypothetical protein |
| **VARPA_3060** | 29.45 | hypothetical protein |
| **VARPA_5951** | 28.71 | dihydrodipicolinate synthetase( EC:4.2.1.52 ) |
| **VARPA_0468** | 28.14 | hypothetical protein |
| **VARPA_1128** | 27.39 | cyanophycin synthetase( EC:6.- ) |
| **VARPA_3061** | 27.12 | hypothetical protein |
| **VARPA_5797** | 26.16 | yiaab two helix domain-containing protein |
| **VARPA_2489** | 24.51 | phosphate ABC transporter, periplasmic phosphate-binding protein |

**Table 2.** Top 25 uniquely downregulated loci in biofilm cultures

| LOCUS | FOLD<br>REPRESSION | GENE ANNOTATION |
| --- | --- | --- |
| VARPA_4525 | 18.64 | extracellular solute-binding protein, family 7 |
| VARPA_1647 | 12.43 | indolepyruvate oxidoreductase subunit b( EC:1.2.7.8 ) |
| VARPA_0430 | 11.76 | sigma 54 modulation protein/ribosomal protein s30ea |
| VARPA_2331 | 10.40 | glycine cleavage system h protein |
| VARPA_2332 | 8.47 | glycine cleavage system t protein( EC:2.1.2.10 ) |
| VARPA_1832 | 7.87 | hypothetical protein |
| VARPA_1660 | 7.78 | extracellular ligand-binding receptor |
| VARPA_4079 | 7.72 | efflux transporter, rnd family, mfp subunit |
| VARPA_5437 | 7.37 | hypothetical protein |
| VARPA_4653 | 7.27 | aminotransferase class-iii( EC:4.1.1.64 ) |
| VARPA_4408 | 6.80 | hypothetical protein |
| VARPA_0874 | 6.79 | hypothetical protein |
| VARPA_3019 | 6.47 | hypothetical protein |
| VARPA_1470 | 6.43 | hypothetical protein |
| VARPA_2596 | 6.34 | hypothetical protein |
| VARPA_5615 | 6.07 | extracellular ligand-binding receptor |
| VARPA_2330 | 6.00 | glycine dehydrogenase( EC:1.4.4.2 ) |
| VARPA_5988 | 5.88 | hypothetical protein |
| VARPA_4455 | 5.85 | hypothetical protein |
| VARPA_4664 | 5.75 | two component transcriptional regulator luxR family |
| VARPA_2423 | 5.74 | hypothetical protein |
| VARPA_1645 | 5.71 | transcriptional regulator marr family |
| VARPA_4080 | 5.70 | outer membrane efflux protein |
| VARPA_2366 | 5.70 | gcn5-related N-acetyltransferase |
| VARPA_3579 | 5.65 | membrane alanyl aminopeptidase( EC:3.4.11.2 ) |

**Figure 5.**
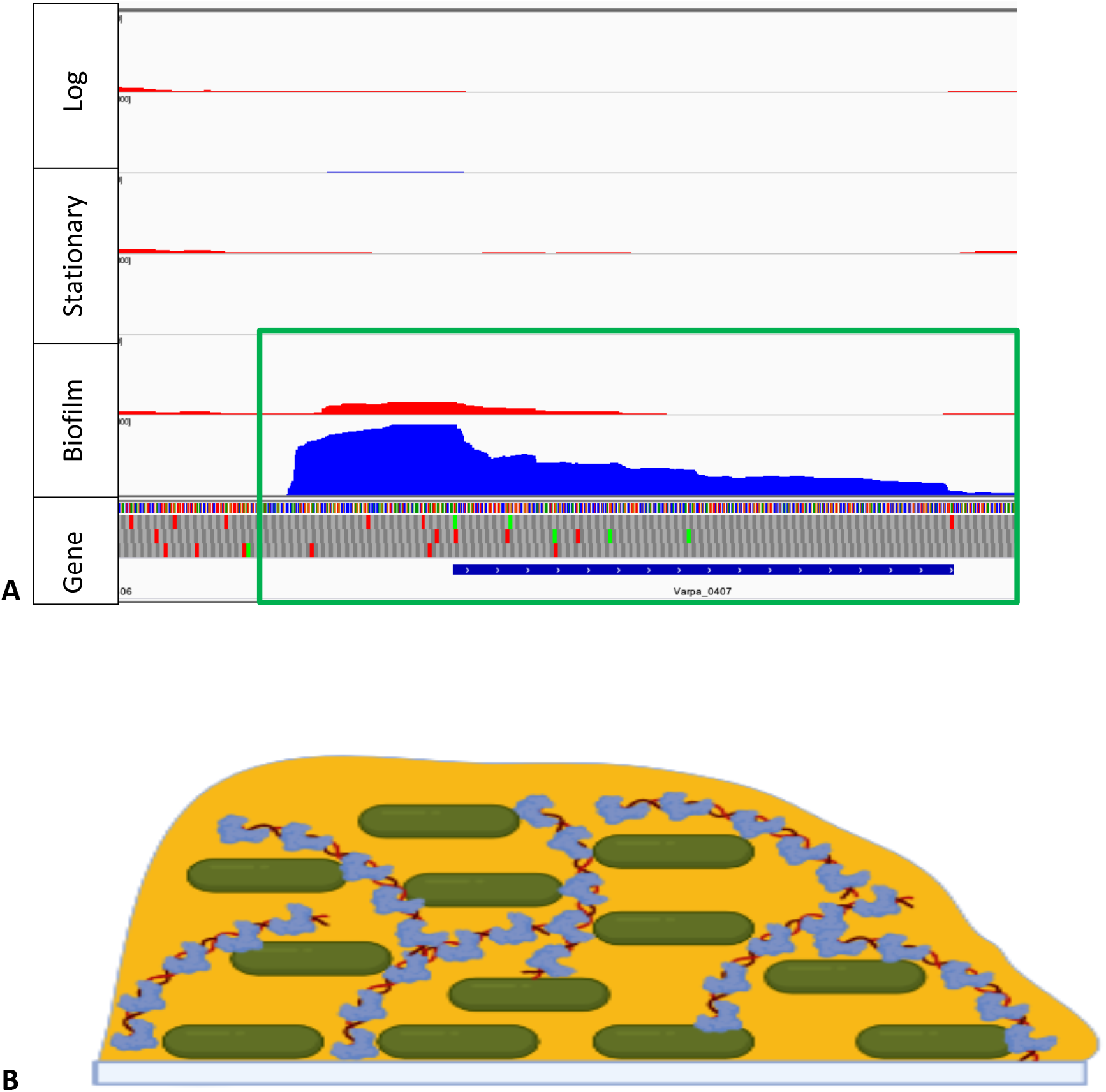
A) Varpa_0407 transcriptome data showing leader region, antisense expression only in biofilm growth (green box). B) model of biofilm eDNA protection by Snf2 superfamily secreted proteins.

### RNA turnover role in Biofilm Regulation

The most specifically highly upregulated gene in our analysis was Varpa_1640 (Table 1, **green**), which was not identified in our previous biofilm mutant screen. This protein is predicted to be a DEAD-box RNA helicase, a widespread family of proteins recently shown to play a role in biofilm formation [26]. We hypothesize that this protein is active in RNA turnover in *V. paradoxus* EPS based on the KEGG predicted RNA degradation pathway [27] (Figure 6A). The RNA degradosomes Type A-C (representing machinery identified in *E. coli, Pseudomonas*, and *Rhodobacter*, respectively) are overlapping but well represented in the *V. paradoxus* EPS genome, consisting of polynucleotide phosphorylase (PNPase, Varpa_4029), enolase (Varpa_2167), RNaseE/R (Varpa_1519, Varpa_3333), and helicases (DEAD/DEAH box, Varpa_4256, Varpa_0178, Varpa_2773), along with the RNA 5’ pyrophosphohydrolase RppH (Varpa_4825) and the Rho transcription termination factor (Varpa_2500). Varpa_1640 was not identified in the KEGG orthology as part of this machinery, nor was the RhlB protein present in the KEGG assignments (Figure 6A) but the STRING network map [28] for Varpa_1640 connects this locus to the RNA degradosome (Figure 6B).The other DEAD/DEAH box helicase (Varpa_4256) is upregulated 15x specifically in biofilms, while none of the other proteins involved in RNA degradation are specifically upregulated in this growth phenotype. The PNPase, enolase, and RNAses are all highly expressed in biofilm (**Supplemental Data S1, S2**), leading to the hypothesis that differential expression of Varpa_1640 is a mechanism for regulating RNA stability and drastically altering the gene expression profile. A relationship between a DEAD-box helicase and biofilm growth was recently described in the plant pathogen *Xanthomonas citri* [26], based on mutations in the HrpB helicase resulting in reduced expression of Type IV pili. Intriguingly, our prior work indicated that Type IV pili are critical for the switch between biofilm and swarming phenotypes in *V. paradoxus* EPS [15], suggesting a possible link between RNA stability, pilus formation, and attachment phenotypes.

**Figure 6.**
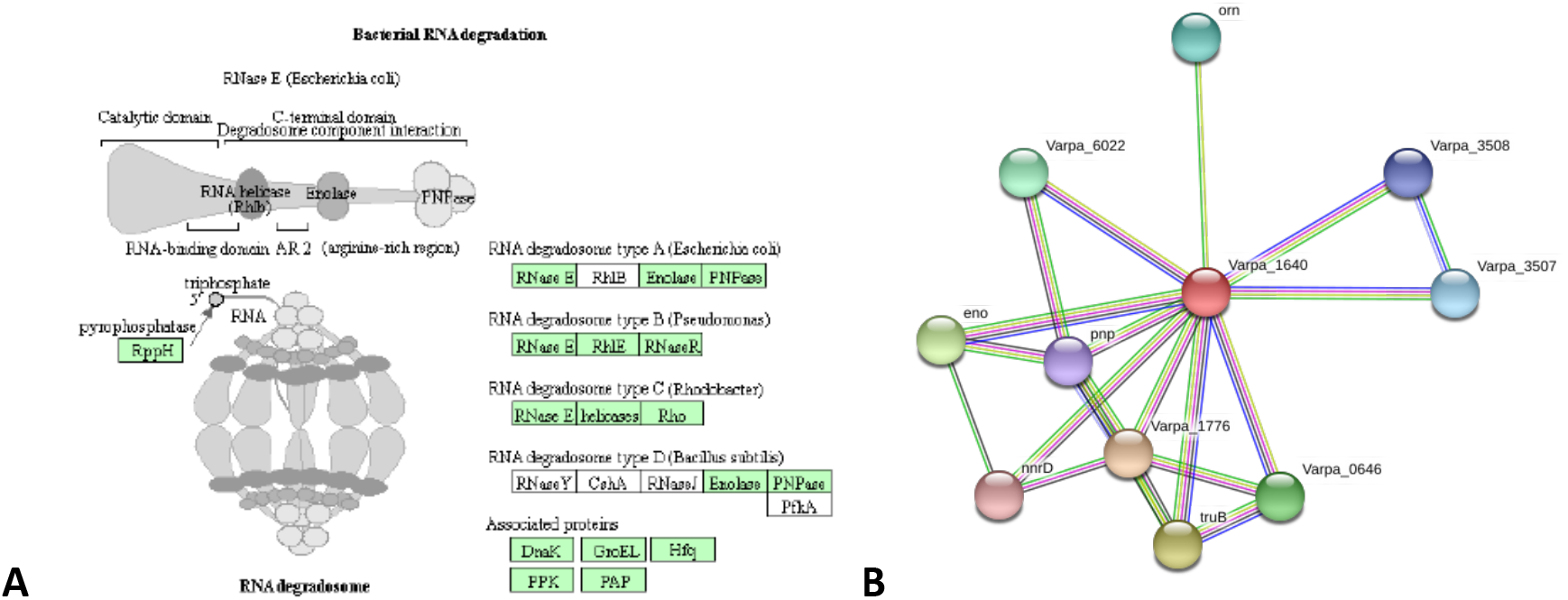
**A)** Proposed role of Varpa_1640 in RNA turnover. Although it is not annotated as such, Varpa_1640 could fit functionally into the type A-C degradosomes in the place of RhlB, RhlE, or unspecified helicases **B)** STRING protein interaction network for Varpa_1640 showing associations with components of RNA degradosome.

### Regulation of Pilus formation and the *tad* locus

The *V. paradoxus* EPS genome encodes multiple pili, including a Type IV pilus with multiple PilY1 proteins and a putative *tad*/*flp* pilus locus (Varpa 5148-5160) similar to the *tad* locus characterized in the bacterium *Aggregatibacter actinomycetemcomitans* [29]. Diverse roles in surface attachment and motility have been identified for pili in different bacterial systems especially type IV pili (T4P) [30], making inference about the roles of these appendages difficult. The tad/flp pilin is classified as a T4Pb class pilin, and is broadly distributed in both gram negative and gram positive bacteria [31]. This pilus has been identified as the appendage responsible for tight adhesion, and also as the bundle forming pilus, and has been implicated in adherence to surfaces and virulence in several different bacteria [32–34]. In *Pseudomonas aeruginosa* pilus gene expression changes in different strains and biofilm conditions can vary widely and be difficult to interpret [2][35]. Inspection of the set of *V. paradoxus* EPS genes with altered expression for pilus related loci revealed that all of the identified pilus genes were downregulated in biofilm with the exception of Varpa_3516 and Varpa_3520. However, the expression levels at these loci (which are predicted to be cotranscribed – see **Supplemental Data S2**) are quite minimal, so it is not clear that this signal is meaningful. The *tad* locus genes are predicted to be expressed in three distinct transcripts (**Supplemental Data S2**), with only the pilus gene and the prepilin peptidase having a very large fold decrease in expression in biofilms (**Supplemental Data S1**). The putative prepilin peptidase (Varpa_5149) was annotated as a pseudogene initially but in more recent annotations has been identified as an open reading frame (not shown, https://www.ncbi.nlm.nih.gov/nuccore/NC_014931.1?report=graph). The putative operons for the rest of the tight adhesion pilus are also downregulated, but to a much lower degree than the pilin and peptidase (Figure 7B). The presence of a strong antisense RNA signal in this operon (Figure 7B, **green box**) suggests potential regulation by RNA stability, as this rise in potential double stranded RNA formation corresponds with the lower transcript levels specifically in biofilm growth. An independent experiment attempting to enrich for motility mutants identified transposon insertions within the *tad* locus leading to deficiencies in swarming motility (Figure 7A). Using a Tn5 insertion disrupting the Varpa_5148 pilin gene, we showed using complementation *in trans* with different constructs (Figure 7A) that both the pilin and the prepilin peptidase (Varpa_5149) are necessary and sufficient to restore motility in this background (Figure 7C). This data supports experimentally the operon structure suggested by transcriptome analysis, and is evidence that this pilus is directly involved in swarming motility, which has not previously been shown in any system. Previously, only *Pseudomonas aeruginosa* had been shown to contain both Type IVa and IVb pili [34], and this is the first time to our knowledge that the tad/flp pilus has been associated directly with motility.

**Figure 7.**
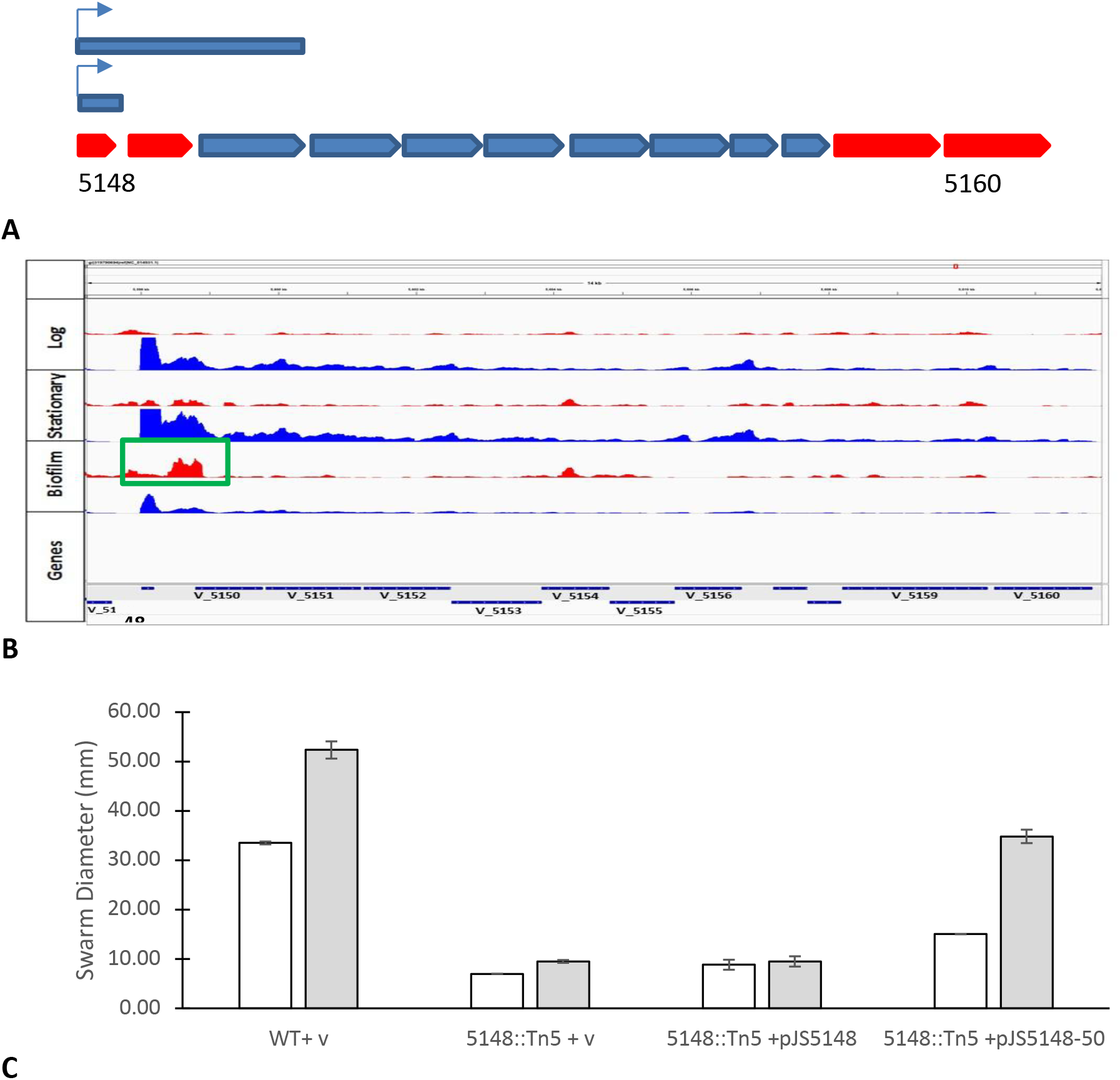
Putative tight adhesion pilus locus map and complementation. **A**) locus map and complementation constructs (blue bars). Red orfs indicate sites where transposon insertions resulted in reduced motility. **B**) representative transcript profiles at the putative tad locus in all three tested phases of growth. A potential antisense transcript (green box) is identified associated specifically with biofilm growth. **C**) Complementation analysis suggests that the putative tad pilus is required for swarming motility. No biofilm phenotype was associated with this mutation.

## CONCLUSIONS

We show here that the rhizosphere isolate *Variovorax paradoxus* EPS has a substantial shift in gene expression when it grows in a biofilm, with about 28% of its genome specifically altered in expression in a static biofilm model. Analysis of the expression pattern has led to a potential new hypothesis about the role of RNA stability in this phenotype, and the specific role of the Varpa_1640 DEAD-box helicase in this mechanism. A previously unrecognized type of small secreted DNA binding protein was identified and is proposed to have an important and specific role in biofilm growth. In contrast with previous work in other systems, the expression profile, presence of a cis-antisense RNA, and mutational analysis suggest that the *flp/tad* locus is suppressed in biofilms and expressed during motile growth. Mutational analysis of biofilm formation yields sharply contrasting results compared to transcriptomic analysis, suggesting both methods are necessary for a complete picture of this complex phenotype.

## ACKNOWLEDGEMENTS

The authors would like to thank NIGMS for funding 1R15GM116173-01 and 1R15GM090242-01, which supported PMO, RJF, and MJP. We would also like to thank NIGMS 1R25GM100829-01A1 MBRS RISE program for supporting JLS.

